# Interrelationships among key Reproductive Health indicators in Sub-Saharan Africa

**DOI:** 10.1101/430207

**Authors:** Mulu Abraha Woldegiorgis, Denny Meyer, Janet E. Hiller, Wubegzier Mekonnen, Jahar Bhowmik

## Abstract

**Introduction:** Indicators of reproductive health (RH) services, outputs, outcomes and impacts are expected to be related with each other and with key social determinants. As the provision of RH services is usually integrated, the effort expended to improve one component is also expected to affect the other components. There is a lack of evidence-based models demonstrating the interrelationships among these indicators and between RH indicators and social determinants.

**Objective:** To examine interrelationships among key RH indicators and their relationship with key social determinants in Sub-Saharan Africa (SSA).

**Method:** This study used data from the most recent demographic and health survey conducted during the period from 2010 to 2016 in 391 provinces of 29 SSA countries. We focused on seven RH indicator — antenatal care, skilled birth attendance, postnatal care, contraceptive prevalence rate (CPR), ideal number of children, birth interval and total fertility rate (TFR), along with selected socio-demographic indicators. The unit of analysis was sub-national, at provincial level. Structural equation modelling was used to examine the strength of interrelationships among the indicators based on the total standardized effect sizes. Significance tests and 95% confidence intervals for the total effects were presented using a bias-corrected bootstrap method.

**Results:** Women’s literacy rate, at the centre of the model, has direct connections with all the RH indicators included in the final model. The strongest relationship was observed between women’s literacy rate and CPR with a total standardized (std.) effect size of 0.79 (95% CI: 0.74, 0.83). RH indicators are interrelated directly and/or indirectly. A strong direct effect was also observed in the relationship between CPR and birth interval (β=0.63, 95%: 0.50, 0.77) and the model suggests that the reported ideal number of children is a key predictor of birth interval (Std. effect size=-0.58, 95% CI: -0.69, -0.48) and TFR (Std. effect size=0.52, 95% CI: 0.38, 0.62).

**Conclusion:** RH indicators are strongly interrelated and are all associated with women’s literacy. The model of interrelationships developed in this study may guide the design, implementation and evaluation of RH policies and programs.

## Introduction

Ideally, every country’s health system is intended to improve the health and wellbeing of its citizens by reducing both morbidity and mortality rates through evidence-based policy, implemented through program management, research, and partnerships with other stakeholders. Health ministries, at different levels of the health system, work in collaboration with different stakeholders to achieve the best from the health service programs guided by an overarching strategy. The provision of integrated Reproductive health (RH) services is a key component of a health care system. The integration is apparent from the top level where a harmonized effort results in coordination of available resources to the grass-roots level where RH services are provided for needy mothers in a one-stop-shop style [1, 2].

Reproductive health (RH), an implied component of the World Health Organization’s definition of health, implies that women and men have the right to be informed of and to have access to the safe, effective, affordable and acceptable methods of fertility regulation of their choice. It includes the right of all women to have access to appropriate, safe and affordable obstetric care, birth control and health care services [3], because it is vital to their lives, their babies, their families and the society at large [4].

Much attention is given to improving RH in developing countries as indicated by the existence of different policies, standard management strategies and guidelines that are put in place by international agencies and governments to improve RH service uptake and to prevent, manage and report simple and complicated maternal cases [5, 6]. Nonetheless, the burden of disease associated with RH in the developing world is still high [7].

Sub-Saharan Africa (SSA), a region with the highest population growth rate in the world [8], is known to be one of the developing regions with the lowest RH status. The maternal mortality ratio, which is considered to be the key impact indicator of health status among all public health statistics, is high in the region, mainly due to high fertility rates [9], low coverage and poor quality of services, along with the influence of challenging operating environments and harmful traditional practices in the region [10].

RH services are best provided in an integrated manner [11]. In a region where the density of skilled health professionals is low [12], the integrated approach has the potential to improve synergy and maximize efficiency [4]. The integration of RH services has many other benefits including improvement of availability of services and supporting staff training that can improve opportunities for early identification and treatment of health problems [13]. Most importantly, it leads to fulfilment of client needs by providing an integrated and client centred service [14]. Accordingly, there are considerable efforts made to integrate RH services in SSA [15, 16].

Family planning, which reduces pregnancy and child birth related risks by increasing birth interval [17], is enhanced by effective maternal health services. These services are provided as a continuum of care from conception to the postnatal period. This usually starts with the diagnosis of pregnancy, followed by provision of antenatal care (ANC) and skilled birth attendance (SBA) services. Postnatal care (PNC) services, which start immediately after delivery, includes another important component of RH – fertility control [18]. During postnatal visits, mothers receive counselling services about family planning (FP) options so that they can make informed decisions about birth control [19]. Moreover, exclusive breast feeding in the first six months after birth, sometime called the lactational amenorrhea period (LAM), provides effective protection against pregnancy [20].

There has been a debate about the impact of FP programs on fertility reduction, focusing on the relative importance of reducing fertility desire and improving access to FP for reducing fertility [21, 22]. It was once stated that the difference in total fertility rate (TFR) across countries can mainly be explained by differences in fertility desire, not by access to contraception [23, 24]. On the other hand, it has been established that FP efforts are important not only for reducing fertility but also for reducing fertility desire through access to information [25, 26].

Besides service integration, RH indicators are thought to be interrelated due to common social, demographic and cultural pre-disposing factors which may affect each of the RH indicators, although the strength of these effects may vary from one indicator to another [27–29].

It is hypothesised that women’s literacy is the most important predictor among all other socio-demographic predictors. A multi-country study in SSA by Bongaarts [30] emphasised the importance of education on decreasing TFR. This is due to combined effects: women’s literacy improves demand for and utilization of contraceptives and also decreases fertility desire. The study also highlighted that differences in the educational attainment of women strengthens the relationship among the RH indicators. Other studies also emphasized the importance of education for improving RH status. Literate women have better health outcomes [31] because they are likely to have better access to information and services [32], and better health service seeking behaviour [33]. With these facts in mind, we hypothesise that women’s literacy rate is the strongest common predictor of RH indicators.

Despite the assumption of interrelationship among the RH indicators, there is limited evidence on the actual level of interrelationship among these indicators in the SSA region. Existing literature has focused on relationships among specific components of RH such as ANC and SBA [34], fertility desire, demand for and utilization of family planning as well as fertility status [30] or the relationship of these variables with key determinants [27]. However, the relative strength of these relationships has not yet been systematically investigated. This study examined the interrelationship among the RH indicators and their association with key social determinants in SSA.

## Conceptual model

In order to establish the extent to which the RH indicators are interrelated, a conceptual model was developed as shown in Figure 1. This approach, sometimes called model generating, produces a tentative and simple initial model based on facts and literature which may be modified after being tested using the actual dataset [35]. After removing a few indicators due to potential multi-collinearity, the model had seven observed RH indicators — antenatal care, skilled birth attendance, postnatal care, contraceptive prevalence rate, ideal number of children, birth interval and total fertility rate, and one socio-demographic indicator — women’s literacy rate. In this model, TFR is the overall outcome indicator whereas women’s literacy rate is assumed to be the main predictor of better RH, with direct relationships to seven of the RH indicators. It is hypothesised that provinces with higher literacy rates tend to increase their mean birth intervals and reduce their mean ideal number of children, leading to lower TFRs. A higher ideal number of children is hypothesised to contribute to shorter birth intervals and subsequent higher TFRs. In addition, it is hypothesised that better PNC will increase the contraception prevalence rate (CPR), which in turn will decrease the TFR.

**Figure 1:**
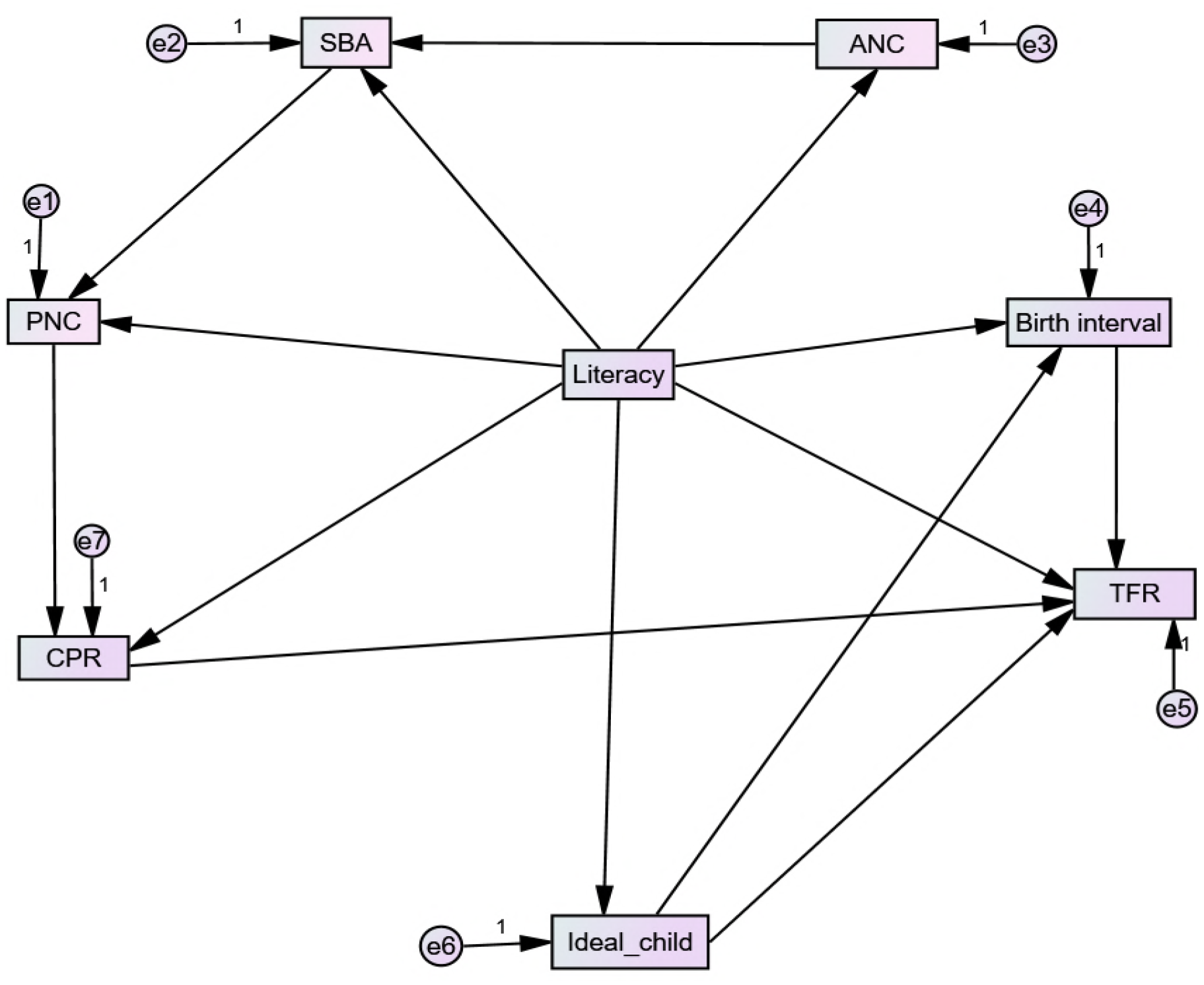
Conceptual framework of interrelationship among reproductive health indicators. ANC=Antenatal care coverage, SBA=skilled birth attendance coverage, PNC= postnatal care coverage, CPR=contraceptive prevalence rate, Ideal child=mean ideal number of children, TFR=total fertility rate, Birth interval= median birth interval, Literacy=women’s literacy rate.

The RH indicators included the conceptual model form four groups, namely maternal health service (ANC, SBA, PNC), fertility control (CPR), fertility desire (ideal number of children) and fertility status (birth interval and TFR). It is also hypothesised that indicators representing each RH group, for instance the three maternal health service indicators, are more related to each other than indicators representing another group (e.g, fertility control).

## Methods

### Description of data source

The study used provincial (regional) key RH and other socio-demographic indicators sourced from the demographic and health survey (DHS) database [36]. The unit of analysis for the indicator data was province (region), a sub-country with definable administrative and geographic characteristics. Initially, a total of 490 provinces were included from the most recent (as of September 2017) standard DHS data of 31 SSA countries with at least one standard DHS between 2010 and 2016. While all DHS data were aggregated at the province level, provinces with missing data for at least one indicator were excluded from the analysis. As shown in Appendix 1, the final sample size was 391 provinces from 29 countries.

### Study indicators

This study used selected RH indicators from each of the four RH thematic areas: maternal health, fertility control, fertility desire, and fertility status.

The indicator data at provincial level were aggregated using means, medians or percentages. Percentage values were taken for coverage data such as ANC, SBA, PNC and CPR. Mean values were taken for TFR and ideal number of children, and the median value was taken for birth interval as per DHS’s aggregation practice [37]. Descriptions of each indicator and method of aggregation (percentage, mean or median) are displayed in Appendix 2.

### Data analysis

Correlational analysis was used to assess the relationship among the RH indicators and the relationships among these indicators and some key socio-demographic factors, including literacy, access to mass media, access to health care, whether wife beating is justified given one or more reasons, whether the woman participates in all household decisions (Appendix 3).

Structural equation modelling (SEM) was applied to test and describe the extent of the relationships between the various indicators using AMOS version 24. SEM is a technique used to describe linear relationships among measured or latent variables [38]. In this particular paper, we applied path analysis to examine the interrelationship among the measured RH indicators. SEM allows the testing of a number of interrelationships simultaneously [39]. It can be used for both exploratory and confirmatory research. This allowed the testing of the initial conceptual model shown in Figure 1 and the subsequent development of a model which better described the data.

### Model assessment, modification and validation

Chi-squared goodness of fit can be used to determine if the hypothesized model describes the data well [40]. However, due to the limitations of chi-square statistics, their sensitivity to sample size and reliance on the assumption of normality [41, 42], we also considered other absolute and relative goodness of fit indices. Root mean square error of approximation (RMSEA) and Probability of Close Fit (PCLOSE = P(RMSEA<5% in population)) were used as absolute fit indices, whereas the Normed-fit index (NFI), Goodness-of-fit statistic (GFI) and Comparative fit index (CFI) were used as relative fit indices. RMSEA less than 0.08, PCLOSE greater than 0.05, GFI, NFI and CFI more than 0.95 are associated with good model fit [40, 42], producing acceptable type I and type II error rates.

The data were randomly split 50:50 into a training and a validation dataset with the training data used to evaluate the initial conceptual model as described above. When the model fit was found to be inadequate, the training data was used to further develop the conceptual model based on theory and using the standardized residual covariances to identify missing paths.

Standardized residual covariances are defined as the difference between the sample covariance and the model-implied covariance for each pair of indicator variables. Regression weights were added one at a time to the model linking the pair of indicators with the highest standardized residual covariance in absolute value. This was done until all standardized residuals were less than two in absolute value. Simultaneously, paths with non-significant regression weights were removed from the model, ensuring that the model was not over-fitted. The validation data was then used to evaluate the goodness of fit for this final model. The standardized coefficients were reported for all paths. Significance tests and 95% confidence intervals for the total effects were calculated using the bias-corrected bootstrap method.

## Results

### Background characteristics (see Appendix 1)

A total of 391 provinces from 29 countries were considered in this study. Among the provinces included in the study, the highest number were in Tanzania and Malawi whereas Comoros contributed the smallest number of provinces. Democratic Republic of Congo, Cote D’viore, Ethiopia, Ghana and Mozambique had 11 provinces, similar to the regional median number of provinces. The regional mean women’s literacy rate was 56.5%. While Lesotho had the highest (97%) literacy rate, the lowest (14%) was observed in Niger. The overall prevalence of teenage pregnancy in the region was 24.3%, the highest being in Niger (40.4%). Less than half of married women in the region (47.1%) agreed with at least one reason supporting the view that wife beating is justified, but this percentage was 76.3% for Malian women.

### Reproductive Health characteristics

Mean values for RH indicators by country are displayed in Table 1. ANC had the highest coverage among the RH indicators in the region. The majority of the countries (62%) had more than 90% ANC coverage while only four countries achieved less than 70% coverage. The median SBA coverage was 64%, and 50% of the countries achieved between 54% and 81% with the median institutional delivery slightly lower (median=63%, IQR=23). PNC coverage was relatively low in almost every country with a median regional coverage of 31% (IQR=32%). The median regional TFR was 5 children, with a minimum of 3.3 in Lesotho to a maximum of 7.6 in Niger. With a regional median coverage of 18%, CPR ranged from 5% to 66%.

**Table 1:** Percentage and mean values of RH indicators in SSA countries between 2010 and 2016.

| Country (Survey period) | ANC (%) | SBA (%) | Inst. Delivery (%) | PNC (%) | CPR (%) | Unmet need (%) | Demand sat (%) | birth interval (median) | ideal child mean) | Wanted TFR (Mean) | TFR (mean) |
| --- | --- | --- | --- | --- | --- | --- | --- | --- | --- | --- | --- |
| Angola (2015/16) | 81.6 | 49.6 | 45.6 | 18.3 | 12.5 | 38.0 | 24.3 | 30.8 | 4.9 | 5.2 | 6.2 |
| Benin (2011/12) | 85.8 | 84.1 | 86.9 | 47.0 | 7.9 | 32.6 | 17.4 | 35.6 | 4.6 | 4.0 | 4.9 |
| Burkina Faso (2010) | 94.9 | 23 | 66.3 | 12.7 | 15.0 | 24.5 | 36.9 | 35.9 | 5.5 | 5.4 | 6.0 |
| Burundi (2010) | 98.9 | 60.3 | 59.5 | 29.4 | 17.7 | 32.4 | 32.7 | 32.0 | 4.2 | 4.5 | 6.4 |
| Cameroon (2011) | 84.7 | 63.6 | 61.2 | 33.1 | 14.4 | 23.5 | 30.8 | 32.7 | 5.5 | 4.5 | 5.1 |
| Chad (2014/15) | 63.7 | 24.3 | 21.7 | 11.6 | 5.0 | 22.9 | 17.6 | 29.3 | 8.2 | 6.1 | 6.4 |
| Comoros (2012) | 92.1 | 82.2 | 76.1 | 17.4 | 14.2 | 32.3 | 27.4 | 31.0 | 5.3 | 3.8 | 4.3 |
| Congo (2011/12) | 85.9 | 83.1 | 91.5 | 54.9 | 20.0 | 18.4 | 31.7 | 38.8 | 5.0 | 4.8 | 5.1 |
| Congo Dem. Rep (2013/14) | 88.4 | 80.1 | 79.9 | 31.3 | 7.8 | 27.7 | 16.3 | 30.4 | 6.1 | 5.7 | 6.6 |
| Cote d'Ivoire (2011/12) | 90.6 | 59.4 | 57.4 | 56.1 | 12.5 | 27.1 | 27.5 | 36.8 | 5.2 | 4.5 | 5.0 |
| Ethiopia (2016) | 62.4 | 27.7 | 26.2 | 15.4 | 35.3 | 22.3 | 60.6 | 34.5 | 4.5 | 3.6 | 4.6 |
| Gabon (2012) | 94.7 | 87.1 | 90.2 | 22.2 | 19.4 | 26.5 | 33.7 | 37.6 | 4.6 | 3.4 | 4.1 |
| Gambia (2013) | 98.9 | 64.0 | 62.6 | 55.2 | 8.1 | 24.9 | 23.8 | 34.2 | 6.0 | 4.9 | 5.6 |
| Ghana (2014) | 97.3 | 73.7 | 73.1 | 68.1 | 22.2 | 29.9 | 39.2 | 39.4 | 4.3 | 3.6 | 4.2 |
| Kenya (2014) | 95.5 | 61.8 | 61.2 | 49.1 | 53.2 | 17.5 | 70.7 | 36.3 | 3.6 | 3.0 | 3.9 |
| Lesotho (2014) | 95.2 | 77.9 | 76.5 | 61.5 | 59.8 | 18.4 | 76.1 | 45.8 | 2.6 | 2.3 | 3.3 |
| Liberia (2013) | 95.9 | 61.1 | 55.8 | 54.7 | 19.1 | 31.1 | 37.2 | 37.4 | 4.8 | 4.2 | 4.7 |
| Malawi (2015/16) | 94.8 | 89.8 | 91.4 | 41.3 | 58.1 | 18.7 | 74.6 | 41.0 | 3.7 | 3.4 | 4.4 |
| Mali (2012/13) | 74.2 | 58.6 | 55.0 | 27.2 | 9.9 | 26.0 | 27.2 | 33.5 | 5.9 | 5.3 | 6.1 |
| Namibia (2013) | 96.6 | 88.2 | 87.4 | 68.0 | 55.3 | 17.5 | 75.0 | 45.1 | 3.2 | 2.9 | 3.6 |
| Niger (2012) | 82.8 | 29.3 | 29.8 | 29.0 | 12.2 | 16.0 | 40.8 | 30.9 | 9.2 | 7.4 | 7.6 |
| Nigeria (2013) | 60.6 | 38.1 | 35.8 | 29.9 | 9.8 | 16.1 | 31.3 | 31.7 | 6.5 | 5.2 | 5.5 |
| Rwanda (2014/15) | 98.0 | 90.7 | 90.7 | 41.1 | 47.5 | 18.9 | 65.8 | 38.5 | 3.4 | 3.1 | 4.2 |
| Sierra Leone (2013) | 97.1 | 59.7 | 54.4 | 41.2 | 15.6 | 25.0 | 37.5 | 36.0 | 4.9 | 4.5 | 4.9 |
| Tanzania (2015/16) | 95.2 | 63.7 | 62.6 | 31.4 | 32.0 | 22.1 | 52.9 | 35.0 | 4.7 | 4.5 | 5.2 |
| Togo (2013/14) | 50.0 | 44.6 | 72.5 | 7.5 | 17.3 | 33.6 | 32.3 | 38.0 | 4.3 | 4.1 | 4.8 |
| Uganda (2011) | 94.9 | 58.0 | 57.4 | 29.9 | 26.0 | 34.3 | 40.5 | 30.2 | 4.8 | 4.7 | 6.2 |
| Zambia (2013/14) | 95.7 | 64.2 | 67.4 | 6.4 | 44.8 | 21.1 | 63.8 | 34.9 | 4.7 | 4.5 | 5.3 |
| Zimbabwe (2015) | 93.3 | 78.1 | 77.0 | 43.1 | 65.8 | 10.4 | 85.2 | 43.7 | 3.9 | 3.6 | 4.0 |

*Int. delivery= institutional delivery. The rest abbreviations are written in full under Figure 1*

### Correlation among indicators

Appendix 3 presents a correlation matrix of background variables and the RH indicators. The majority of the background and socio-economic indicators had a significant relationship with each of the RH indicators, however, women’s literacy rate had the strongest correlation with each of the RH indicators. As shown in the table, the strongest correlation was observed between literacy rate and CPR (r=0.79).

Table 2 shows that maternal health indicators were strongly correlated to each other at provincial level. ANC and SBA had a moderate strength correlation (r=0.62). A slightly stronger correlation was observed between SBA and PNC (r=0.66). Despite the expectation of a strong relationship between PNC and CPR, we found only a moderate relationship (r=0.45). On the other hand, there was a weak, negative, linear relationship (r=-0.21) between PNC and unmet need for family planning. As expected, CPR was negatively correlated with unmet need for family planning (r=-0.43). This supports the argument that when women obtain access to contraceptive services, unmet need for family planning decreases.

A strong, positive, linear relationship was observed between mean ideal number of children and TFR (r=0.72). Provinces with higher mean ideal number of children tended to have higher TFR. TFR also had a strong negative correlation with birth interval. A shorter birth interval allows the possibility of having more children, which in turn increases the TFR. There was, therefore, a strong negative relationship between ideal number of children and birth interval. All the bivariate correlations were statistically significant. Most of the significant correlations were moderate to strong in strength, demonstrating substantial associations between the RH indicators.

**Table 2:** Bivariate Correlations among RH indicators.

|  | ANC | SBA | PNC | CPR | TFR | Ideal-child | Birth interval |
| --- | --- | --- | --- | --- | --- | --- | --- |
| ANC | 1 |  |  |  |  |  |  |
| SBA | .618** | 1 |  |  |  |  |  |
| PNC | .500** | .660** | 1 |  |  |  |  |
| CPR | .449** | .653** | .448** | 1 |  |  |  |
| TFR | -.349** | -.639** | -.557** | -.651** | 1 |  |  |
| Ideal-child | -.551** | -.706** | -.505** | -.720** | .721** | 1 |  |
| Birth interval | .408** | .579** | .557** | .724** | -.755** | -.681** | 1 |

\*\*Correlation is significant at the 0.01 level (2-tailed). *Abbreviations are written in full under Figure 1.*

### Relationship of demographic indicators with RH indicators

In line with the previous section – bivariate correlations, the multivariate analysis of covariance (MANCOVA) indicated that socio-demographic indicators had a significant relationship with the combined set of RH indicators. Women’s literacy rate had the strongest relationship (Pillai’s Trace =0.39) followed by access to health care (Pillai’s Trace =0.21).

### Structural Equation Modelling

As shown in Table 3, the conceptual model evaluated by the training data was found to provide a poor fit with the data in terms of both the absolute and relative indices (RMSEA=0.260, NFI=0.825, GFI=0.793 and CFI=0.833). The model was improved progressively, as indicated by the Standardized Residual Covariances, by adding one regression weight at a time as shown in Table 3. Additions were made only when they could be justified in theoretical terms. For instance, it was assumed that provinces with high TFR tend to have low maternal health service coverage. After removing the non-significant regression weights, the final model described the data reasonably well with all the indices within the expected range. The chi-square test was also not statistically significant, confirming an adequate model *(*^2^(8) = 9.15,*p* = 0.330;64.5% *chance of true RMSEA* < 5%).

**Table 3:**
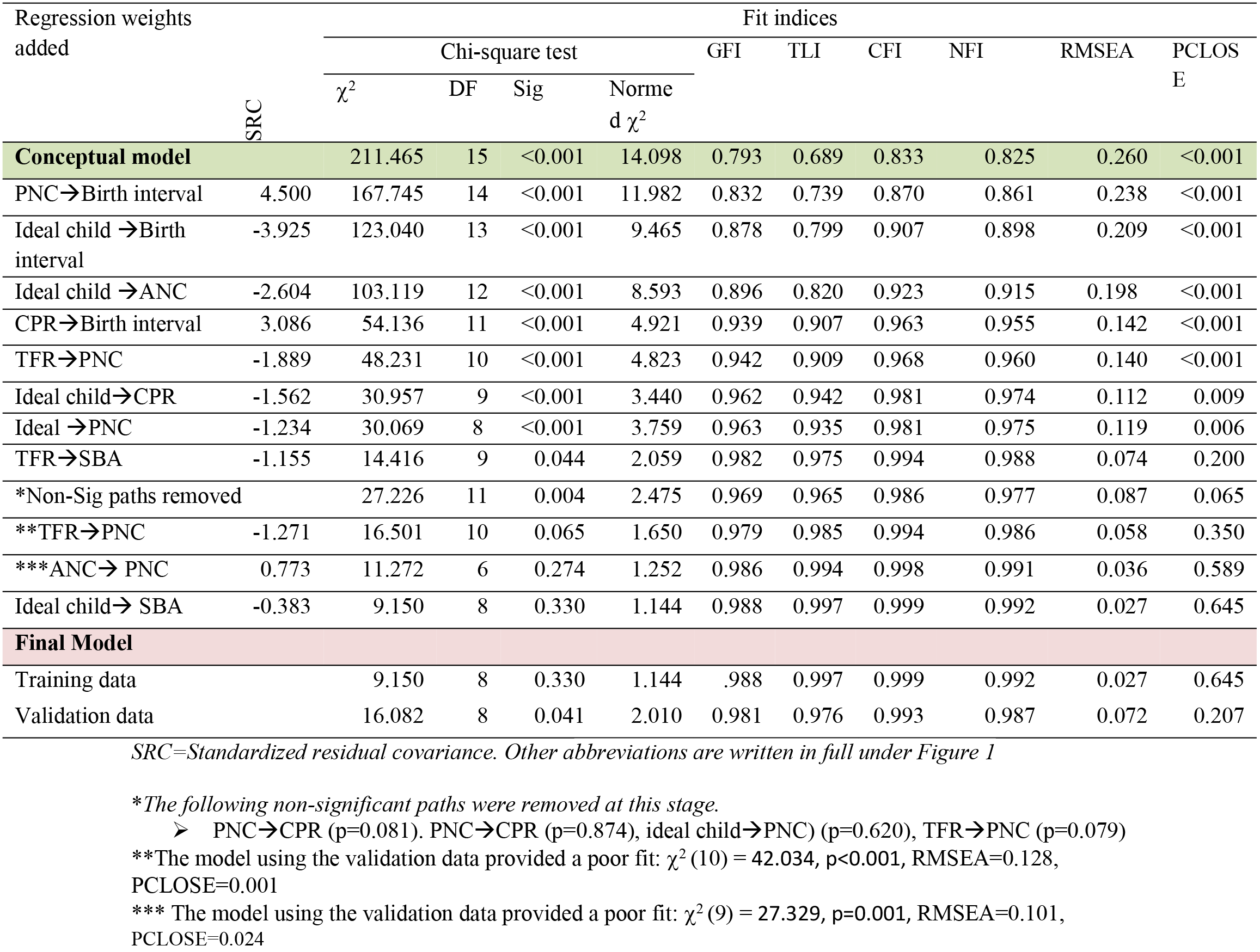
Progressive model fit indices based on standardized residual covariance between indicators.

### Standardized direct effects

The Structural Equation Model in Figure 2 shows the final model with the standardized weights estimating standardized changes in each dependent indicator for a one standard deviation (SD) increase in each independent indicator, when the other independent variables were statistically controlled. These are referred to as standardized (std.) beta coefficients (β). As indicated in this model, women’s literacy was found to be the key predictor having a direct relationship with all seven RH indicators in the final model. For instance, women’s literacy rate was a strong predictor of mean ideal number of children (β=0.70,395% CI: -0.75, -0.63), when controlling for the other RH indicators. However, contrary to expectation, women’s literacy rate had a negative direct relationship with PNC (β=-0.23, 95% CI: -0.37, -0.03) and birth interval (β=-0.26, 95% CI: -0.37, -0.14).

**Figure 2:**
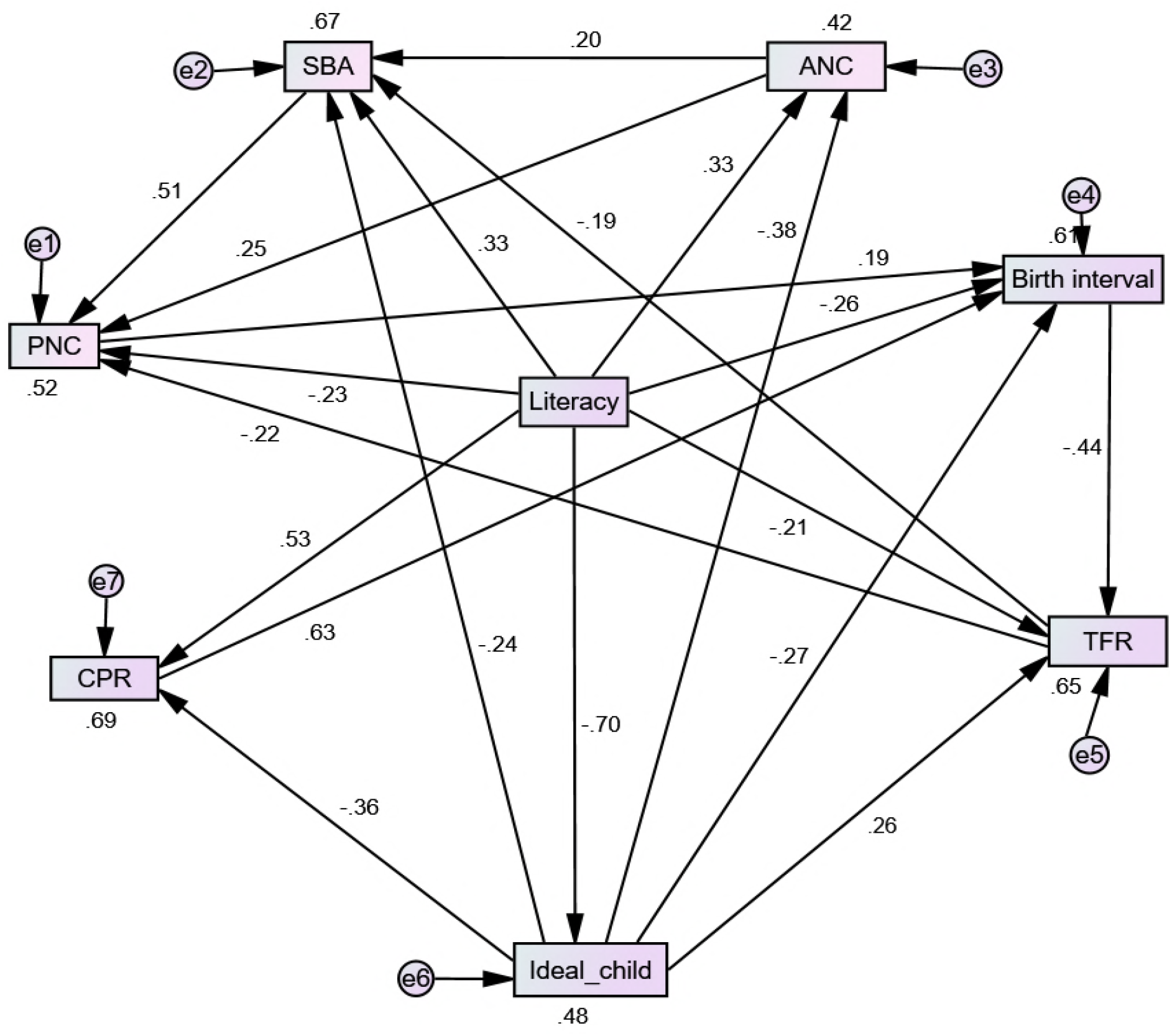
Structural equation model of reproductive health indicators with standardized (β) weights and R-Square values shown. \**Abbreviations are written in full under Figure 1*.

As expected, the RH indicators were interrelated. The strongest direct relationship was observed between CPR and median birth interval β=0.63, 95%CI: 0.50, 0.77). Maternal health indicators were interrelated and were affected by different RH indicators. For a one SD increase in provincial SBA coverage, PNC coverage increased by 0.51 SDs (95% CI: 0.34, 0.63). Provinces with high TFR tended to have low PNC service coverage (β =0.22, CI: -0.36, 0.09). This model explains 65% of the variation in TFR, 69% of the variation in CPR and 67% of the variation in SBA.

### Standardized indirect and total effects

The standardized total effects - the sum of direct and indirect effects - along with the bias-corrected 95% CI for these estimates is presented in Table 4. In the total standardized effects, the strongest relationship was observed between literacy rate and CPR (std. effect size = 0.79 (95% CI: 0.74, 0.83), followed by the relationship between literacy rate and SBA (std. effect size = 0.73 (95% CI: 0.66, 0.78). After taking indirect effects into account, there was a positive relationship between literacy rate and birth interval (Std. effect size = 0.51, 95% CI: 0.42, 0.58). and a positive relationship between literacy rate and PNC (Std. effect size = 0.43, 95% CI: 0.33, 0.53) as originally expected. In addition, there was a negative relationship between TFR and PNC (Std. effect size = -0.32, 95% CI: -0.46, -0.16) and between birth interval and TFR (Std. effect size = -0.45, 95% CI: -0.57, -0.34) as was originally expected. Provinces with lower TFR tended to have higher use of maternal health services, and CPR was a positive predictor of birth interval (Std. effect size=0.65, 95% CI: 0.51, 0.78). Women’s literacy rate was the strongest predictor of TFR (Std. effect size=-0.62, 95% CI: -0.70, -0.54), with higher TFR when literacy rates were low, but the effect of the mean estimate for the ideal number of children was also strong (Std. effect size=0.52, 95% CI: 0.38, 0.62), with a larger mean ideal number of children associated with a higher TFR.

**Table 4:**
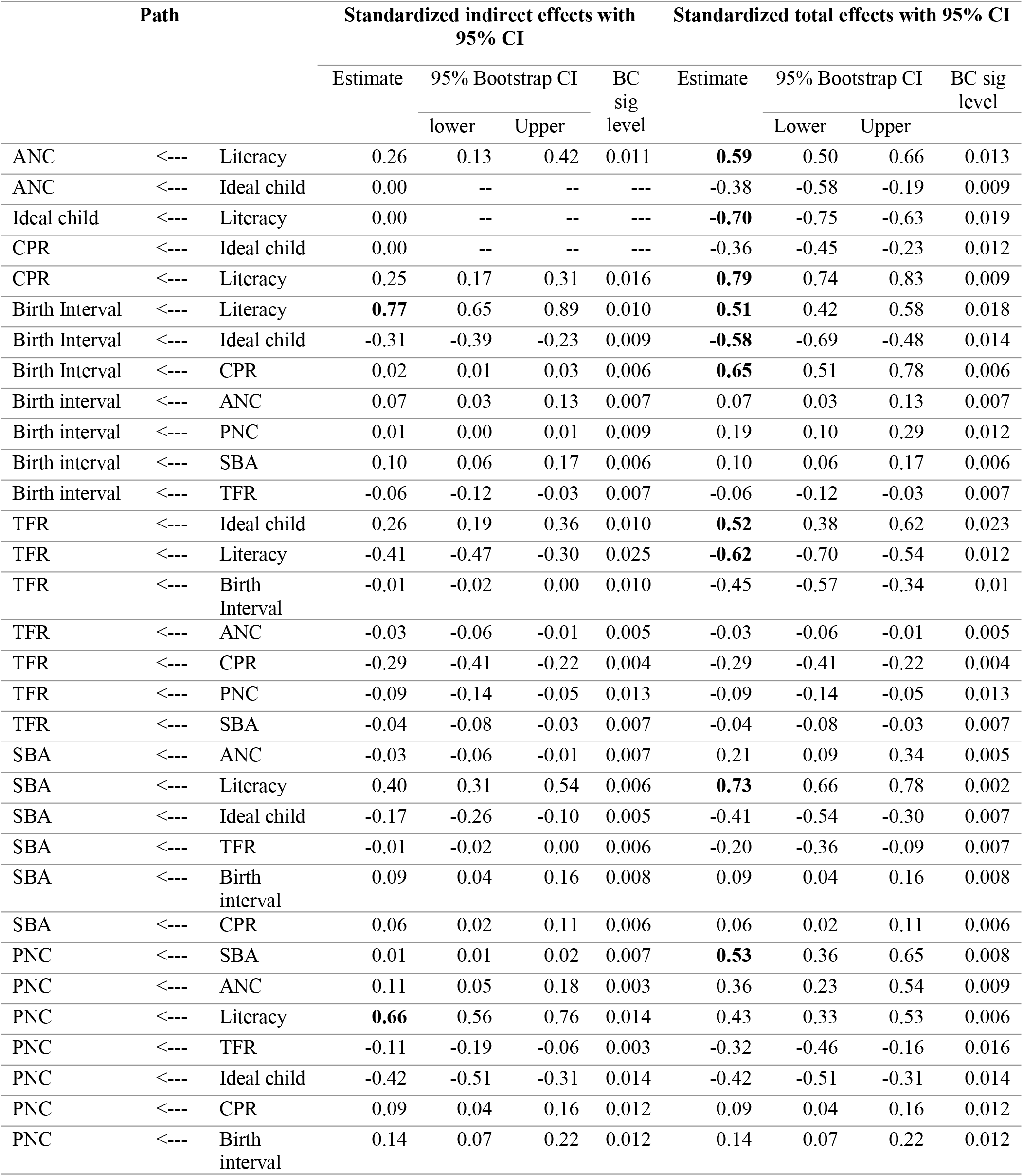
Standardized indirect and total effect sizes with bootstrap generated confidence intervals using the bias-corrected (BC) percentile method (standardized effect size estimates above 0.5 in bold).

## Discussion

This study examined the interrelationship among key RH indicators and selected social determinants. The analysis was conducted at sub-national level where each country in the region was represented by its provinces. The background and socio-demographic indicators had significant relationships with each of the RH indicators. It was found that, in the final model, literacy rate had direct links to all the RH indicators. In line with the existing literature about social determinants of health, especially the power of literacy on the overall health literacy [44], findings in this study suggest that improving women’s literacy rate is a key tool for improving RH service uptake. Moreover, the indirect relationships between women’s literacy and RH indicators implies that improving women’s literacy rate may also have a long-term effect on the key RH status indicators such as TFR.

There were statistically significant relationships among the RH indicators included in this study. This could be due to common background factors such as socio-economic status, educational status, and access to and quality of health care services. It could also be due to the policy landscape of a country and the implementation capacity of the respective provinces. Furthermore, the interrelationships could also be attributed to the integrated provision of RH services within each province. As stated elsewhere [24], neighbourhood cultural effects may also contribute to high fertility desire. Women living in areas where TFR is high or where having more children is the norm, would be less likely to make use of family planning. Ecological interventions, on intervening community and societal factors, to increase awareness and the ability of women to have a manageable number of children may be a key means for improving demand and utilization of family planning services.

The correlation and standardized beta coefficients in the final model suggested that provinces with high ANC service coverage tended to have high SBA coverage. High SBA coverage in turn resulted in increased PNC service uptake. However, the relationships between these indicators were not strong enough for a latent variable called maternal health to be constructed. This indicates that the relationships among these maternal health indicators was not uniform across provinces. Firstly, ANC service addresses service coverage (quantity) rather than service quality [45] which may result in an inflated count in some provinces [46]. Secondly, SBA and PNC services require clinical care services [47] while ANC services can be provided by community health workers in provinces where health facilities are not within travelling distance. Thirdly, there may be differences in the cascade of maternal services from the ANC to PNC across different provinces. This is evidenced by changes in the gap between the coverage of ANC and SBA services among countries across time [27, 28]. These three maternal health services should be treated separately rather than as a single latent variable due to the differential effects of literacy on these services. The effect of literacy on PNC was relatively low (Std. effect size = 0.43) compared to the effects of literacy on ANC (Std. effect size = 0.59) and SBA (Std. effect size = 0.73) in particular.

The strongest association was observed between women’s literacy rate and CPR followed by the relationship between women’s literacy rate and SBA. This suggests that women’s literacy is a key mechanism in order to improve utilization of contraceptive and maternal health services. The standardized total effect sizes indicate that the mean ideal number of children was a negative predictor of CPR as well as median birth interval. However, it was a positive predictor of TFR. At the individual level, a high ideal number of children suggests low demand for contraception. Therefore, it is unlikely there would be ‘*unmet* or *met need’* for family planning; rather it suggests there is ‘*no unmet need’* for family planning. Targeting women with a high ideal number of children should be a priority for family planning services. It was once said that the difficult task is not about reducing unwanted fertility but it is about reducing people’s fertility desire [48].

While improving women’s literacy rate was found to be a key predictor for reducing the ideal number of children, desired number of children was an important predictor of contraceptive use (β=-0.36) indicating a direct pathway. While literacy and ideal number of children had direct and indirect effects on TFR, CPR had an indirect effect on TFR through birth interval. However, the relationship may not be simple as it may be mediated or moderated by important demographic transitions such as change in infant mortality rate, as well as the epidemiological transitions – such as changes in disease patterns. Two theories have been suggested for explaining the positive relationship between TFR and infant mortality [49]. A high TFR may be a response to high infant mortality. On the other hand, it has also been argued that close spacing of births, a key predictor for TFR in the current study, contributes to high infant mortality. In the absence of definitive information about fertility desire and fertility status, previous research suggests that among the key factors driving fertility are fecundability, survival chances, contraceptive practices and other RH milestone practices such as age at first marriage [50]. In the SSA region, where age at first marriage is low [51] and mean CPR is less than 18%, the last two of these factors are expected to play a pivotal role.

Two unexpected findings from the fitted model were the negative direct relationships for literacy rate with PNC and birth interval. These relationships contradicted the simple correlations where a positive relationship was observed in both cases. However, as shown in the model, positive relationships were observed in the total effect sizes for this model, due to the indirect effects through ANC, SBA and CPR services. As expected, therefore, higher literacy rate for a province is associated with higher PNC coverage and longer median birth intervals.

CPR had a strong direct effect on birth interval (Std. effect size =0.65) and birth interval was an important predictor of TFR (Std. effect size =-0.45). These results confirm findings from previous studies about the role of CPR in increasing birth interval and ultimately reducing fertility rates [52]. The negative relationship between TFR and maternal health indicators such as SBA and PNC can partly be explained by the fact that provinces with high TFR tend to have low women’s empowerment, as indicated in this study and elsewhere [53], which negatively impacts on the utilization of maternal health services. Supporting the direction of effect from the TFR to SBA, prior research which examined the effect of parity on SBA services [54], noted that high parity was a negative predictor of SBA service uptake.

The existence of strong relationships among RH indicators suggests that integrated policies and strategies may be more effective in improving RH outcomes than those that focus on a single indicator. Our findings confirm that improving women’s literacy will be critical in improving CPR and thereby reducing fertility in SSA countries with implications for integrated policy making, implementation and evaluation in addressing maternal health and RH issues in the region. More specifically, the model of interrelationships developed in this study may be an important tool in policy analysis as it describes the strength of interrelationships among key RH indicators which are also outcomes/impact indicators of common RH programs in SSA.

The findings of this study also substantiate the need for integrated approaches to maternal health and family planning services in SSA. The fact that women’s literacy came out as a strong predictor of RH indicators suggests that responsibilities for improving RH outcomes is not limited to the healthcare system, but it extends to the educational sectors of countries. Given the considerable interrelationships among RH indicators, system wide effects of programs should be considered in the design, implementation and evaluation of maternal health and population control initiatives and policies in SSA. The model developed in this study would be useful in guiding any evaluation of RH programs in SSA.

This study has demonstrated the application of SEM in examining the interrelationships among RH indicators in SSA. Unlike the traditional regression analysis approaches, SEM provides a basis for a more holistic analysis of interrelationships among the RH indicators. Future studies could consider this approach for analysis of interrelationships in individual level data for specific countries. The model developed in this study could also be used to guide the design and analysis of further research concerning RH indicator relationships using similar techniques.

There are some limitations associated with this study. First, the study focused on aggregated indicator data at province level and there is a possibility of ecological bias [55], because the provinces are heterogeneous even within a country. Second, there existed time differences in the data collection period of DHS from 2010 to 2016 and this could affect the comparability of the data across provinces.

## Conclusions

Based on the findings of this study, the following conclusions were drawn:

1. The SEM developed in this study described the training and the validation data well.
2. In the resulting model, literacy rate stood out as a strong predictor of most of the RH indicators included in this study. It had a direct effect on all RH indicators included in the model. It was also the most important predictor of CPR, ideal number of children, birth interval, SBA, ANC and TFR when indirect and direct effects were considered simultaneously.
3. There existed moderate to high correlation among the RH indicators suggesting that RH indicators are interrelated to each other in the SSA context.
4. Maternal health indicators (especially SBA and PNC) and birth interval were influenced by the majority of RH indicators; and CPR was found to be the main predictor of birth interval.
5. Ideal number of children had both direct and indirect relationship with TFR.
6. The findings of this study confirm the importance of enhancing integrated approaches especially between the education and health sectors.

## Appendix 1: Background characteristics of the women surveyed

| Country | DHS period | *Number of provinces | Women who are literate | Final say in all of the decisions | Access to electricity | Worked in the last 12 months | Teenagers childbearing | Problems in accessing health | Wife beating justified | No access to mass media |
| --- | --- | --- | --- | --- | --- | --- | --- | --- | --- | --- |
| Angola | 2015-16 | 18 | 58.1 | 65.4 | 41.6 | 65.1 | 34.5 | 70.2 | 25.2 | 26.2 |
| Benin | 2011-12 | 18 | 35.1 | 48.3 | 38.4 | 63.6 | 16.5 | 68.6 | 16.2 | 45.7 |
| Burkina Faso | 2010 | 14 | 22.5 | 12.0 | 13.1 | 74.8 | 23.6 | 78.5 | 43.5 | 48.1 |
| Burundi | 2010 | 5 | 61.5 | 49.6 | 5.3 | 73.4 | 9.6 | 86.0 | 72.9 | 38.7 |
| Cameroon | 2011 | 17 | 69.2 | 23.7 | 53.7 | 62.2 | 25.2 | 39.0 | 46.5 | 40.1 |
| Chad | 2014-15 | 21 | 22.1 | 17.4 | 7.7 | 42.9 | 35.9 | 29.3 | 73.5 | 81.4 |
| Comoros | 2012 | 3 | 63.3 | 34.6 | 69.3 | 36.6 | 11.3 | 78.2 | 39.0 | 38.0 |
| Congo | 2011-12 | 14 | 82.2 | 28.4 | 41.6 | 64.2 | 32.9 | 72.8 | 60.7 | 34.8 |
| Congo Dem. Republic | 2013-14 | 11 | 63.8 | 33.5 | 13.5 | 67.2 | 27.2 | 76.0 | 74.8 | 66.3 |
| Cote d'Ivoire | 2011-12 | 11 | 37.7 | 23.6 | 55.8 | 67.1 | 29.6 | 77.6 | 47.9 | 45.5 |
| Ethiopia | 2016 | 11 | 42.0 | 70.6 | 25.6 | 33.3 | 12.5 | 70.0 | 63.0 | 73.6 |
| Gabon | 2012 | 14 | 88.7 | 43.7 | 89.3 | 43.7 | 27.6 | 80.8 | 50.2 | 7.5 |
| Gambia | 2013 | 8 | 45.0 | 39.2 | 44.5 | 42.6 | 17.5 | 43.0 | 58.4 | 29.6 |
| Ghana | 2014 | 11 | 67.1 | 61.6 | 78.3 | 73.4 | 14.2 | 50.7 | 28.3 | 30.8 |
| Kenya | 2014 | 8 | 87.8 | 55.8 | 36.0 | 61.4 | 18.1 | 46.1 | 41.8 | 22.9 |
| Lesotho | 2014 | 10 | 97.0 | 65.4 | 27.8 | 37.8 | 19.1 | 41.8 | 33.3 | 31.7 |
| Liberia | 2013 | 20 | 47.9 | 65.9 | 9.8 | 54.9 | 31.3 | 62.3 | 42.5 | 55.5 |
| Malawi | 2015-16 | 38 | 72.1 | 46.9 | 10.8 | 62.6 | 29.0 | 73.9 | 16.3 | 62.8 |
| Mali | 2012-13 | 9 | 20.6 | 8.9 | 25.6 | 42.6 | 39.3 | 58.6 | 76.3 | 45.8 |
| Namibia | 2013 | 17 | 93.4 | 74.6 | 47.4 | 42.7 | 18.6 | 43.3 | 28.2 | 27.1 |
| Niger | 2012 | 10 | 14.0 | 12.3 | 14.4 | 24.5 | 40.4 | 70.2 | 59.6 | 59.0 |
| Nigeria | 2013 | 6 | 53.1 | 31.3 | 55.6 | 61.8 | 22.5 | 51.0 | 34.7 | 50.4 |
| Rwanda | 2014-15 | 5 | 80.2 | 65.4 | 22.8 | 77.7 | 7.3 | 58.6 | 41.4 | 36.2 |
| Sierra Leone | 2013 | 18 | 35.5 | 45.4 | 13.5 | 68.1 | 27.9 | 71.9 | 62.8 | 56.2 |
| Tanzania | 2015-16 | 39 | 76.8 | 35.2 | 22.5 | 72.3 | 26.7 | 65.5 | 58.0 | 46.1 |
| Togo | 2013-14 | 7 | 52.3 | 29.4 | 45.7 | 70.3 | 16.5 | 65.7 | 28.7 | 39.4 |
| Uganda | 2011 | 8 | 64.2 | 37.5 | 14.6 | 69.3 | 23.8 | 64.9 | 58.3 | 21.0 |
| Zambia | 2013-14 | 10 | 67.5 | 53.8 | 27.9 | 48.8 | 28.5 | 50.2 | 46.9 | 34.1 |
| Zimbabwe | 2015 | 10 | 94.4 | 72.1 | 33.7 | 41.3 | 21.6 | 58.6 | 38.7 | 45.2 |

**\*Total number of provinces =391**
**Wife beating:** Attitude towards wife beating for at least their following reasons: *burns food, argues with husband, goes out without telling husband, neglects children, refuses to have sex with husband,*
**All decisions:** Own health care, making large purchase, visit family, relates and friends
**Access to health:** Problem in any of the following issues: *knowing where to go for treatment, getting permission to go to treatment, getting money for treatment, distance to health facility, having to take transport, not wanting to go alone, concern that there may not be female privier.*

## Appendix 2: Description of reproductive health indicators

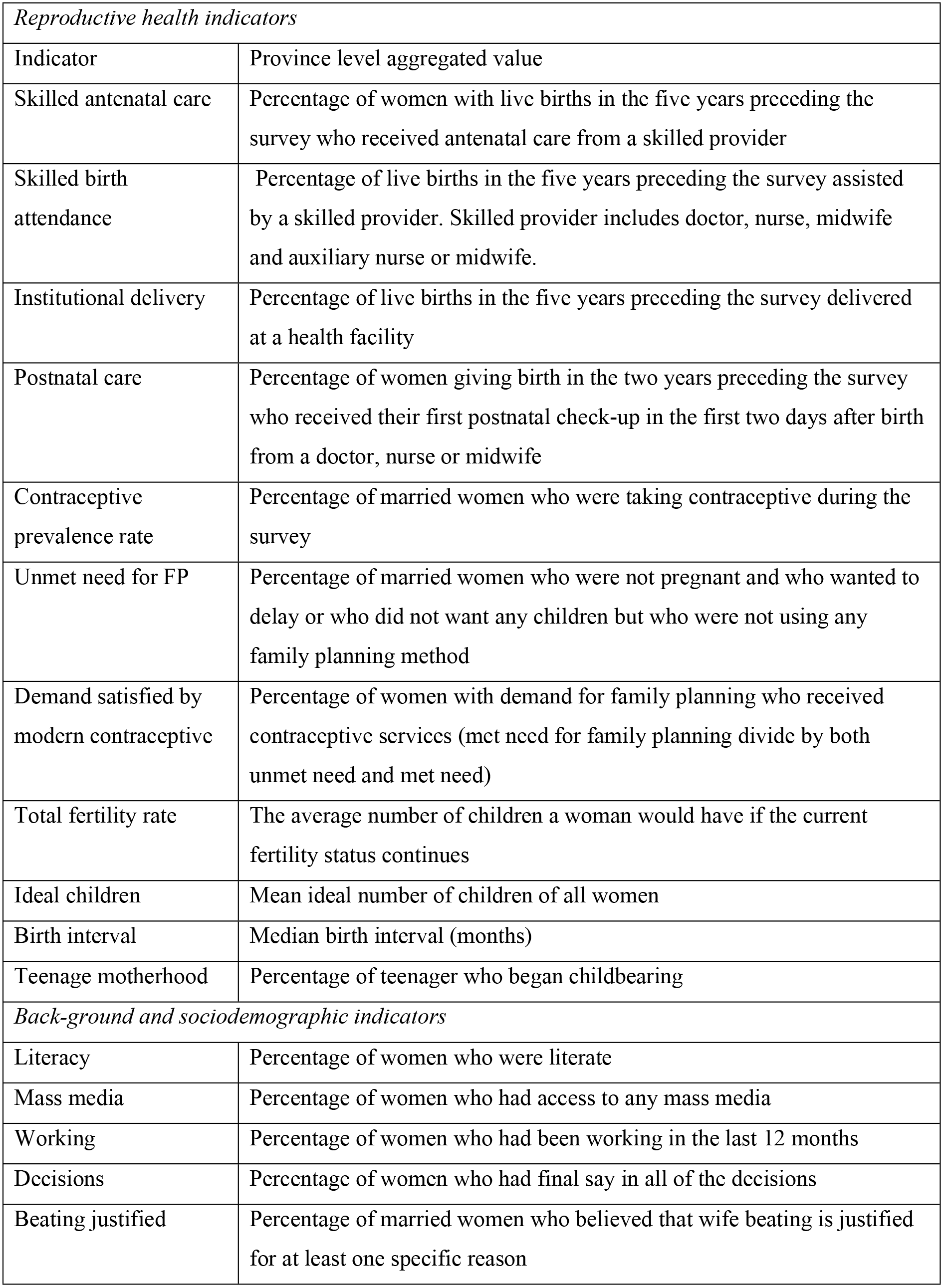

## Appendix 3: Correlation between background indicators with reproductive health indicators

|  | Literacy | Access<br>Mass<br>Media | Access<br>Health<br>Care | Wife<br>beating<br>justified | Final say in<br>all of the<br>decisions |
| --- | --- | --- | --- | --- | --- |
| Antenatal care coverage | .510** | .448** | -0.084 | -.158** | .308** |
| Skilled birth attendance | .732** | .485** | 0.091 | -.432** | .395** |
| Postnatal care coverage | .435** | .314** | .220** | -.322** | .411** |
| Contraceptive prevalence rate | .788** | .301** | .111* | -.365** | .424** |
| Unmet need for family<br>planning | -.299** | 0.075 | -.232** | 0.043 | -0.030 |
| Family planning demand<br>satisfied with contraceptive | .627** | .225** | .127* | -.361** | .448** |
| Total Fertility rate | -.623** | -.442** | -.335** | .478** | -.441** |
| Wanted total fertility rate | -.677** | -.484** | -.244** | .512** | -.500** |
| Mean ideal number of children | -.704** | -.514** | 0.009 | .495** | -.574** |
| Median birth interval | .541** | .313** | .130* | -.430** | .392** |

\*\* . Correlation is significant at the 0.01 level, \* . Correlation is significant at the 0.05 level (2-tailed).

